# Predicted Versus Observed Activity of PCB Mixtures Toward the Ryanodine Receptor

**DOI:** 10.1101/2023.08.22.554299

**Authors:** Justin A. Griffin, Xueshu Li, Hans-Joachim Lehmler, Erika B. Holland

**Author notes:** Address Correspondence to E.B. Holland, California State University of Long Beach, 1250 Bellflower Blvd, Long Beach, CA. 90840.

## Abstract

Non-dioxin-like polychlorinated biphenyls (NDL PCBs) alter the activity of the ryanodine receptor (RyR), and this activity is linked to developmental neurotoxicity. Most work to date has focused on the activity of single congeners rather than relevant mixtures. The current study assessed the RyR activity of single congeners or binary, tertiary, and complex PCB mixtures. Observed mixture activity was then compared to the expected activity calculated using the concentration addition (CA) model or a RyR-specific neurotoxic equivalency scheme (rNEQ). The predictions of the CA model were consistent with the observed activity of binary mixtures at the lower portion of the concentration-response curve, supporting the additivity of RyR1 active PCBs. Findings also show that minimally active congeners can compete for the RyR1 binding site, and congeners that do not activate the RyR1 do not interfere with the activity of a full agonist. Complex PCB mixtures that mimic PCB profiles detected in indoor air, fish tissue, and the serum of mothers and children activated the RyR1 and displayed similar efficacy and potency regardless of varying congener profiles. Neither the CA model nor the rNEQ perfectly predicted the observed activity of complex mixtures, but predictions were often within one magnitude of change from the observed response. Importantly, PCB mixtures approximating profiles found in environmental samples or human serum displayed RyR1 activity at concentrations reported in published research. The work presented will aid in the development of risk assessment platforms for NDL PCBs, and similar compounds, towards RyR1 activation and related neurotoxicity.

## 1. INTRODUCTION

Ryanodine receptors, namely RyR isoforms 1, 2, and 3 (RyR1, 2, or 3) are a family of Ca^2+^ release channels found throughout nervous systems (Pessah *et al*., 2010; Berridge *et al*., 2003). RyR isoforms are embedded in the ER membrane of neuronal cells, including those from the cerebellum, hippocampus, basal ganglia, and cerebral cortex (Abu-Omar et al., 2018). Both RyR1 and RyR2 can be found in neuronal tissue but were first recognized for their important physiological role in skeletal (RyR1) and cardiac (RyR2) muscle health (Pessah et al., 2010). In adults, RyR3 is mainly located in the caudal cortical plate and hippocampus (Pessah et al., 2010). The role the RyR isoforms play in cellular physiology varies greatly by cell type but can contribute to vesicle release, neuronal network development, axon and dendritic growth and movement, and neuronal connectivity (Abu-Omar et al., 2018; Berridge et al., 2003; Del Prete et al., 2014). Additionally, the presence of RyR2 and RyR3 in hippocampal neurons contributes to spatial memory, synaptic plasticity, and long-term memory (Adasme et al., 2011).

The environmental pollutant class, polychlorinated biphenyls (PCBs), alter RyR1 and RyR2 activity (Pessah et al., 2019, 2010). They are man-made organic chemicals produced and utilized in the United States until their production was phased out in the 1970s due to human and wildlife health concerns. There are 209 PCB congeners, but they were produced and used as mixtures in lubricants and coolants found in electrical equipment and as plasticizers in paint, plastics, rubbers, pigments, and dyes (Faroon and Ruiz, 2016). They are stable, highly persistent compounds (Faroon and Ruiz, 2016), such that even after discontinuing their production, they remain in environmental media, including air (Herkert et al., 2018; Marek et al., 2017), water (Chakraborty et al., 2014; Lombard et al., 2023), sediment (Lu et al. 2021), and human (Li et al., 2022; Marek et al., 2013) and wildlife samples (Stahl et al., 2009). Thus, humans are exposed to PCBs through diet (Ampleman et al., 2015; Hardell et al., 2010) and inhalation (Herkert et al., 2018; Marek et al., 2017), the latter of which has been gaining increased attention regarding PCB levels detected in schools (Hamper, 2020; Marek et al., 2017; Osterberg and Scammell, 2016). Following exposure, PCBs distribute into lipid-rich tissues of the body (Hardell *et al*., 2010) and can cross the blood-placenta barrier during pregnancy and be transferred from mother to child through lactation (van den Berg et al., 2017). Studies have shown that PCBs can accumulate in fetal (Lanting et al., 1998), infant, and adult brains showing complex concentration differences across age, brain region, and sex (Chu et al., 2003; Dewailly et al., 1999; Li et al., 2022). Additionally, during body weight loss, PCBs mobilize from lipid-rich tissue into the blood with risk of redistribution to other organs such as the brain (Brown et al., 2019; Fonnum and Mariussen, 2009).

Out of the 209 congeners, there are 197 PCBs, often referred to as non-dioxin-like PCBs (NDL PCBs), many of which have been shown or are predicted to activate RyR channels (Holland et al., 2017a; Pessah et al., 2006; Rayne and Forest, 2010). The NDL PCBs represent more than 60% of the total PCB burden found in environmental and human samples (Marek et al., 2017, 2013; Stahl et al., 2009), and exposure to NDL PCBs have been linked to neurotoxic outcomes (reviewed by Klocke et al. 2020; Pessah et al. 2019). Briefly, previous work with isolated RyR protein preparations, primary cells, cell lines, and animal models show that NDL PCB activity at the RyR channel altered intracellular Ca^2+^ homeostasis (Wayman et al., 2012), neuronal growth (Wayman et al., 2012), and muscular physiology (Niknam et al., 2013). Ca^2+^ dyshomeostasis, especially during neuronal development, may alter neural networks and synaptic plasticity important to neuronal behavior, such as memory and spontaneous activity (Pessah et al., 2019). Supporting the role of NDL PCBs in neurodevelopmental toxicity include studies by Yang et al. (2009) showing that weanling rats exposed to Aroclor 1254, a commercial mixture containing high amounts of NDL PCBs, experienced alerted dendritic growth and deficits in learning and memory. It was also found that PCB 95, a tri-ortho NDL PCB that is highly potent and efficacious at the RyR (Pessah et al., 2006), alters neuronal structure such as dendritic growth. These findings were not observed with PCB 66, which displays minimal RyR activity (Pessah et al., 2006). These observations and results by other groups (reviewed by Klocke and Lein, 2020; Pessah et al., 2019) identify NDL PCBs as an environmental risk factor for neurodevelopmental toxicity.

To date, research has mainly focused on assessing the activity or toxicity of single congeners (Holland et al., 2017a; Pessah et al., 2006; Sethi et al., 2019). The current study investigated the RyR1 activity of NDL PCBs as single congeners and binary and tertiary mixtures containing congeners with varying potency and efficacy. Additionally, the study assessed the activity of complex PCB mixtures that mimic congener profiles found in indoor air (Marek et al., 2017), fish tissue (Stahl et al., 2009), and human serum (Marek et al., 2013). The activity of the binary, tertiary, and complex mixtures was then compared to the expected RyR1 activity calculated using the Concentration Additional Model (CA model; Faust et al. 2000; Jonker et al. 2005; Belden and Lydy 2006; Kirakosyan et al. 2009; Fritsch and Pessah 2013) or a RyR1-specific neurotoxic equivalency scheme (rNEQ; Holland and Pessah 2021). Together this work provides an insight into the activity of complex PCB mixtures toward RyR1 channels, which are important to neuronal signaling and other physiological pathways. Moreover, our results aid in the development of risk assessment platforms for environmentally relevant NDL PCBs mixtures towards RyR1 activation and neurotoxic risks.

## 2. MATERIALS AND METHODS

### 2.1 Individual PCBs and Mixtures

Individual PCB congeners were purchased neat from AccuStandard (99.5–100% pure; New Haven, CT, USA) and dissolved in dry DMSO. To create binary mixtures, congeners were combined in a 1 : 1 ratio in a test tube. Binary mixtures combined the RyR1 full agonist (PCB 95) with another full agonist (PCB 149), a partial agonist (PCB 52), a minimally active congener (PCB 47), or an inactive congener (PCB 126). The tertiary mixture was created to represent new findings regarding non-Aroclor PCBs (PCB 47, 51, and 68) detected in the indoor air of new or remolded buildings (Table S1; Herkert et al. 2018).

The RyR1 binding activity of complex mixtures created to mimic the PCB congener profile from environmental and human samples was also assessed. Complex PCB mixtures found in the environmental or human serum (Tables S2-S9) were custom prepared and the final PCB mixtures characterized by GC-MS/MS as described in the Supplementary Data. Created mixtures included the 25 congeners with the greatest contribution to the total PCBs reported in published works (Marek et al., 2017, 2013; Stahl et al., 2009). Some samples did not have 25 congeners detected above the limit of quantification (LOQ) in which case only those that appeared above the LOQ were included in the mixture creation and mixture toxicity assessment. Overall, eight environmentally relevant mixtures were created, including PCB mixtures found in the indoor air of urban and rural schools (Tables S2 and S3), the tissue of predatory and bottom-dwelling fish from Cook County, IL (Tables S4 and S5), and the serum from mothers and children living in urban or rural locations (Tables S6-S9). The PCB concentrations that were reported in the indoor air of middle and high schools (Marek et al., 2017) and the concentrations found in the serum of mothers and children (Marek et al., 2013) were part of a large AESOP (Airborne Exposures to Semi-volatile Organic Pollutants) cohort study investigating exposures in mothers and children from East Chicago, IN or Columbus Junction, IA. Fish data were part of a National Lake Fish Residue study which detected all 209 PCB congeners in predatory and bottom-dwelling fish from US lakes (Stahl et al., 2009), and raw data by lake were provided by Dr. Leanne Stahl (USEPA Office of Water; Washington, DC).

### 2.2 Radioactive Binding Assay

The junctional sarcoplasmic reticulum (JSR), containing concentrated RyR1, was a generous gift from Dr. Isaac Pessah and Dr. Feng Wei (University of California Davis) and was created as described previously (Pessah et al., 2006) using the skeletal muscle from a male New Zealand white rabbit Binding assays were performed as reported previously (Holland et al., 2017a) in assays consisting of 0.5 mL binding buffer (140 mM KCl, 15 mM NaCl, 20 mM HEPES, pH 7.2) with 24 μg/mL JSR protein in the presence of 50 μM CaCl, 1 nM [^3^H] Ryanodine (56 Ci/mmol; Perkin Elmer Life and Analytical Sciences, Waltham, MA, USA). Treatments included a 0.5% DMSO (v/v) vehicle control or single PCBs, binary, tertiary, or complex PCB mixtures in the presence of 0.5% DMSO. PCB concentrations in the assay ranged from 0.01-50 µM. Non-specific binding assays were run under the same condition in the presence of 10 μM ryanodine and 200 μM EGTA. Tubes were incubated for 3 h at 37 °C and assays were terminated using three washes of ice-cold harvesting buffer (140 mM KCl, 15 mM NaCl, 20 mM HEPES, pH 7.2) and vacuum filtered onto a GF/B filter (Whatman, Clifton NJ, USA). Filters were soaked in liquid scintillation fluid (BD Cocktail, Fisher Scientific, Waltham, MA, USA), held overnight, and then tritium levels determined on a Beckman Coulter LS 6500. Specific binding was determined by subtracting the non-specific binding from the total binding.

### 2.3 Development of Concentration Response Curves

Experimentally observed concentration-response curves (CRC) for the RyR1 activity of single congeners or binary, tertiary, and complex mixtures were developed using a four-parameter non-linear regression curve in Prism 7.0 (Graphpad Software, San Diego, CA) and graphed onto semi-log plots. Sigmoidal curves were used to determine effective concentration (EC) values, such as the EC_50_ and EC_2X_ (concentration needed to cause a 200% overactivation), and maximal responses to describe potency and efficacy, respectively. The absolute EC_50_ values for single congeners and mixtures were calculated by normalizing response curves to the maximal response observed for PCB 95; a highly potent and efficacious congener (Holland et al., 2017a; Pessah et al., 2006) with extensive data available for toxicity toward the RyR and related physiological pathways (Pessah et al., 2019).

### 2.4 Comparison to Predictive Models

Observed full CRCs or EC values for mixtures were compared to expected curves or EC values developed using the CA model. The CA model is used to examine the additivity of mixtures (Belden and Lydy, 2006; Faust et al., 2000; Fritsch and Pessah, 2013; Jonker et al., 2005; Kirakosyan et al., 2009) and states that chemical mixtures display additivity when their combined effects do not deviate from an expected response calculated from the following equation (Eq. 1):

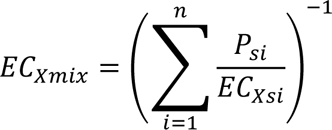

where EC*_xmix_* is the PCB concentration of the mixture that is expected to cause an activation effect (e.g., EC_50_), P_si_ is the proportion of the single PCB of interest present in the mixture and EC*_Xsi_* is the concentration of the PCB of interest shown to cause an activation effect (e.g., EC_50_). Values ranging from the EC_2.5_ - EC_97.5_ of single PCB congers were used to develop an expected curve for binary or tertiary mixtures using the CA model (see Eq. 1) following that described previously (Fritsch et al. 2013). A Goodness of fit test was then used to compare the CA model expected CRC to the observed CRC. Expected EC values were also compared to the 95% confidence intervals (CI) for the corresponding observed EC values.

The tertiary and environmental mixtures created (see Table S1-S9) varied slightly between the actual congener profile found in published work (Herkert et al., 2018; Marek et al., 2017, 2013; Stahl et al., 2009). Expected response values developed with the CA model were calculated using the created congener profile for comparison to the observed EC values gathered in binding assays. Previous work (Holland et al., 2017a; Pessah et al., 2006) reported relative EC_50_ values, rather than absolute EC_50_ values, which are not appropriate for use in mixture assessments (Wagner et al., 2013). Thus, previously published or predicted EC_2X_ values for a given congener (Holland et al., 2017a; Pessah et al., 2006; Rayne and Forest, 2010) were used for inclusion in Eq. 1. The CA model expected EC_2X_ was then compared to the EC_2X_ of the observed EC_2X_ gathered from the CRC for the corresponding mixture.

Y-interpolation of the observed response curves for complex mixtures (Tables S2-S9; Prism 7.0) was used to evaluate the RyR activity of actual concentrations detected in environmental or serum PCB mixtures. Additionally, the previously published PCB 202 response curve (Holland et al., 2017a) was used to perform Y-interpolation of PCB 202 bioassay equivalent concentrations developed using a RyR-specific neurotoxicity scheme (rNEQ) described previously (Holland and Pessah, 2021). Here, EC_2X_ values of each congener were used to develop bioassay relative potency (REP) values relative to the EC_2X_ of PCB 202, the most potent RyR1 active congener tested to date (Holland et al., 2017a). Congener REPs were then multiplied by concentrations a given PCB in a mixture and summed to calculate a PCB 202 equivalent for a given mixture. The rNEQ bioassay equivalent concentration was then used to predict RyR activity observed on the CRC of PCB 202 (Holland et al., 2017a).

## RESULTS

### Single Congeners

Ryanodine receptor binding data for individual congeners can be found in Table 1, and CRCs can be found in accompanying figures with binary or tertiary mixture data (Figure 1-4). The well-studied PCB 95 was highly potent and efficacious, in line with previously published data (Pessah et al. 2006 and Holland et al. 2017). Compared to PCB 95, PCB 149 displayed a slightly lower maximal effect but was within the 95% confidence interval of the PCB 95 maximal effect. Therefore, PCB 149 was considered a full agonist for use in binary mixtures. PCB 52 was less potent and less efficacious than PCB 95 and was considered a partial agonist. Finally, the para-substituted PCB 126 was inactive at the RyR1, consistent with previous work (Pessah et al. 2006). PCB 47, 51, and 68, which lacked previous RyR activity data, all activated the receptor but only PCB 51 caused appreciable efficacy with a maximum effect of 800% (Table 1). These congeners have recently been shown to represent a large percentage of the total PCBs detected in the air of newly constructed or remodeled homes (Herkert et al., 2018).

**Table 1.**
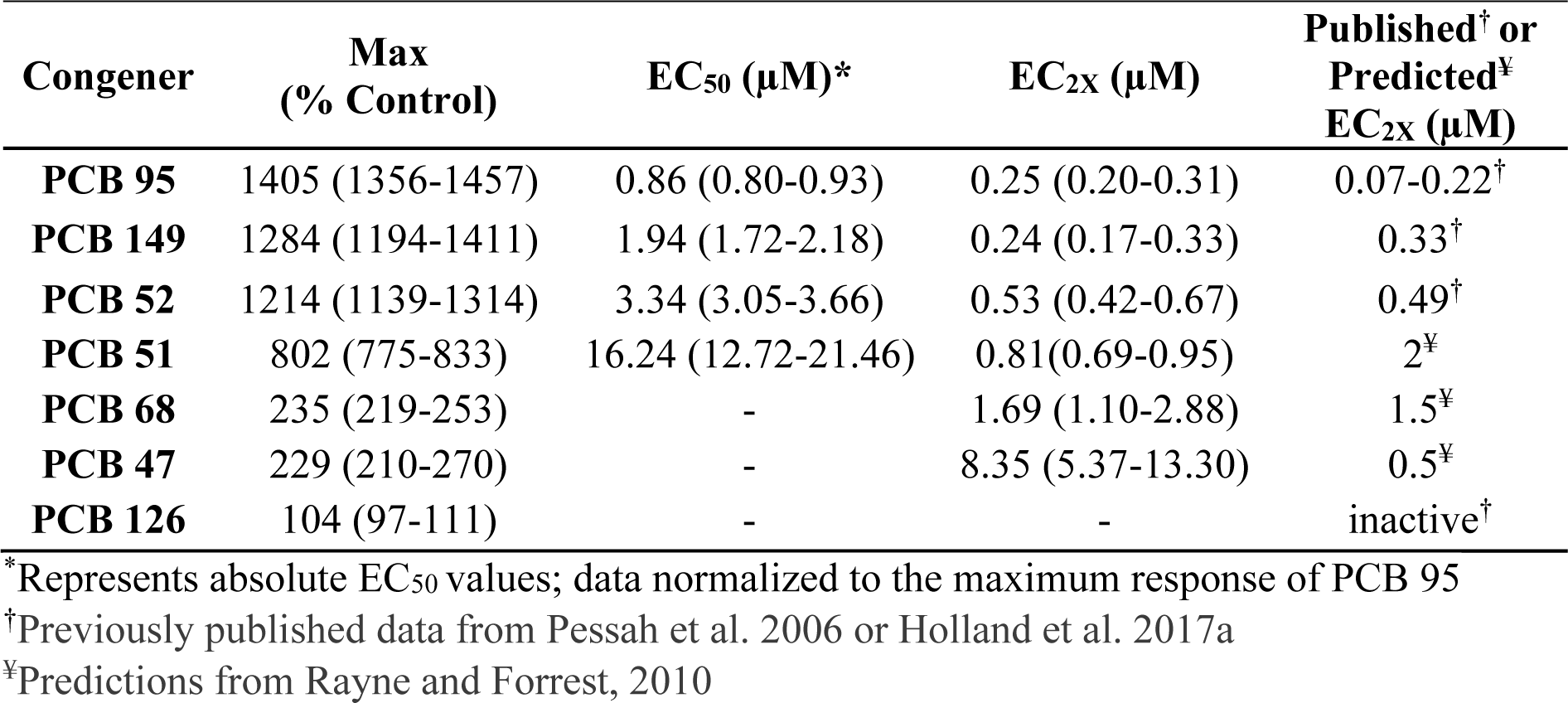
Single Congener Potency and Efficacy Toward the Ryanodine Receptor, Isoform 1, in Rabbit Skeletal Muscle. Values in parenthesis represent the 95% Confidence Interval for a given parameter.

**Figure 1.**
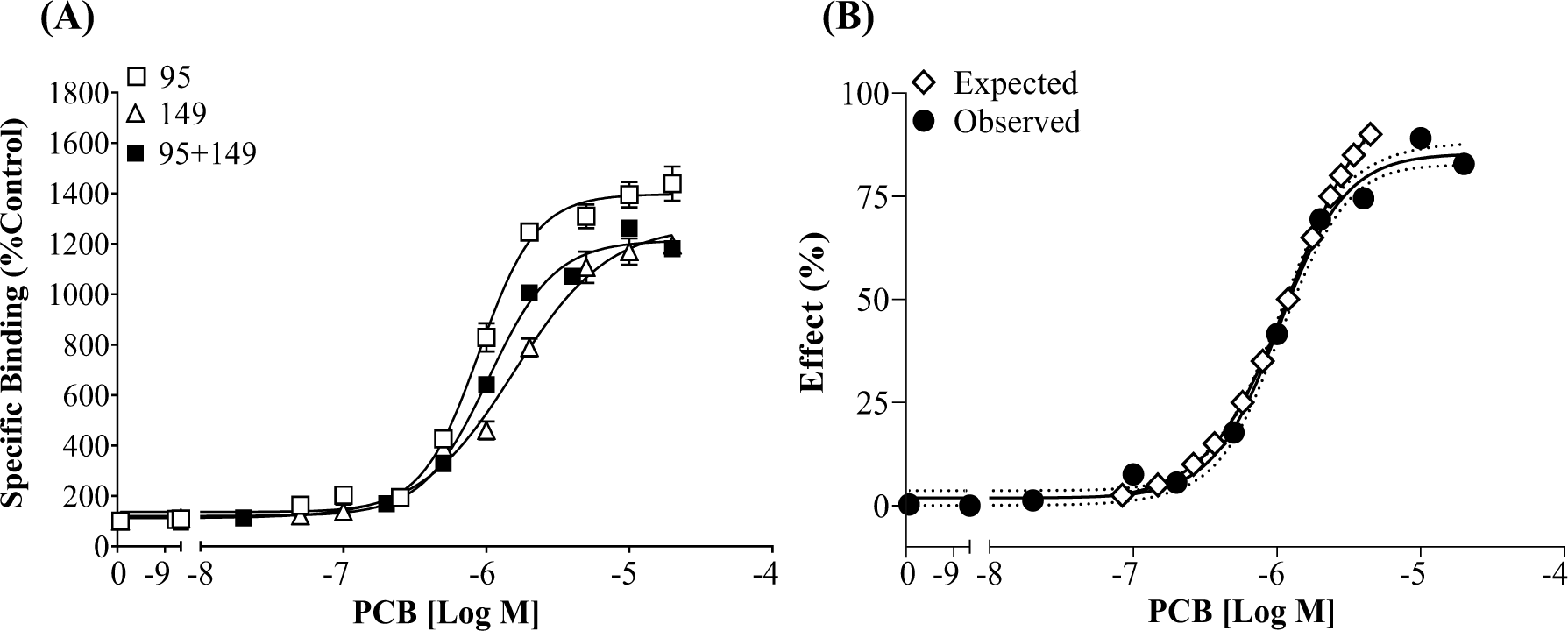
Full agonists PCB 95 and PCB 149 display additivity toward the RyR1 when present in a binary mixture. (A) [^3^H] Ry specific binding is displayed as a percentage relative to the DMSO control; (B) specific binding of the mixture is normalized with a baseline set at 0 and a maximal response set at 100% and compared to the CA model expected curve. Dotted lines represent the 95% CI for the observed response curve. Values represent Means ± SEM. PCB 95 (n=12), PCB 149 (n=9), mixture (n=6).

### Binary and Tertiary Mixtures

When the two full agonists, PCB 95 and PCB 149, were combined, the observed curve resulted in a maximum response of 1220% compared to the DMSO control and had an EC_2X_ of 0.27 μM (Figure 1A). This is surprising given that PCB 95 and PCB 149 displayed higher efficacy when ran alone. If the full range of EC values (EC_2.5_ – EC_97.5_) were included, the CA model expected curve was significantly different from the observed CRC for the PCB 95 + PCB 149 mixture (Figure 1B; F=5.88, p < 0.01). This discrepancy was primarily driven by differences in the expected efficacy and the observed efficacy. If concentrations expected to cause >85% effect in the CA model were removed, the expected effect curve did not vary significantly from the observed effect curve (F = 0.97; p = 0.43). Additionally, the ECs for the lower portion of the curves overlapped. For example, the expected EC_25_ aligned with that observed in the binary mixture curve (Exp: EC_25_ = 0.58 μM; Obs: EC_25_ = 0. 53 μM, 95% CI 0.46-0.61 μM). However, there was a minor difference between the EC_50_ values (Exp: EC_50_ = 1.19 μM; Obs: EC_50_ = 1.39 μM, 95%CI 1.26-1.54 μM). Together these data suggest that PCB 95 and PCB 149 display additivity; however, the predictions of the CA model are not accurate for higher EC values, given the slight difference in the efficacy observed between the congeners.

When PCB 95 and PCB 52 were combined, the binary mixture led to a maximum response of 1290% compared to the DMSO control and had an EC_2X_ of 0.27 μM (95% CI 0.21-0.35: Figure 2A). When the observed curve was normalized to the PCB 95 maximum response (i.e.,100%), it showed an approximate 85% activation. The CA model expected curve was not developed for the PCB 95 and PCB 52 binary mixture effects above 80% of the maximal response for RyR1. This was because, as a partial agonist, those values could not be calculated from the single congener curve for PCB 52. The expected effect curve developed using the CA model did not vary significantly from the observed effect curve (Figure 2B; F = 1.33; p = 0.27). The ECs overlapped between the observed and expected curves on the lower portion. For example, the expected EC_25_ did not vary from the EC_25_ of the observed curve (Exp: EC_25_ = 0.72 μM; Obs: EC_25_ = 0.73 μM, 95% CI 0.59-0.82 μM). Similar to the PCB 95 + PCB 149 curve, the EC_50_ values did display a minor difference, where the predicted value fell slightly outside of the 95% CI of the observed EC_50_ (Exp: EC_50_ = 1.37 μM; Obs: EC_50_ = 1.58 μM, 95%CI 1.44-1.73 μM,). Again, this suggests that active congeners act in an additive manner, but the CA model cannot accurately predict the higher effect concentrations of congeners with varying efficacy.

**Figure 2.**
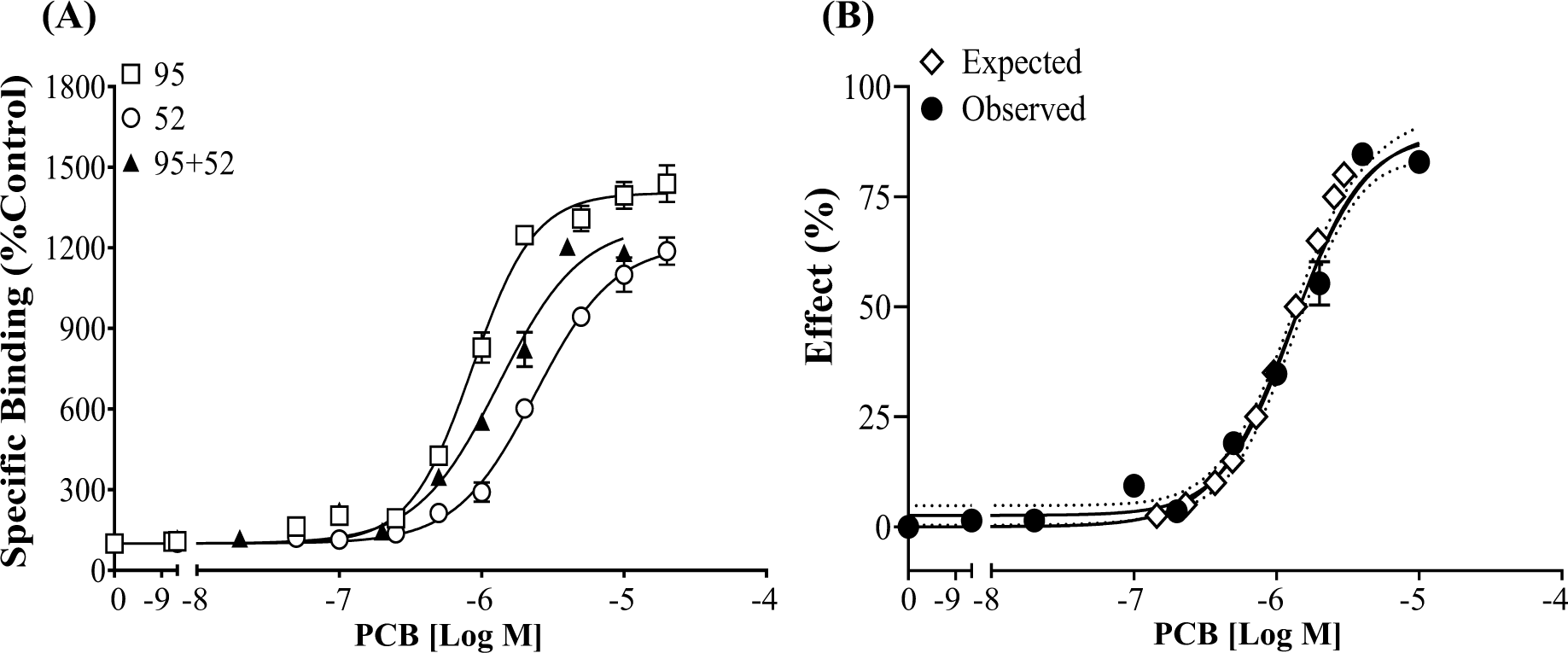
Full agonist PCB 95 and partial agonist PCB 52 display additivity toward the RyR1 when present in a binary mixture. (A) [^3^H] Ry specific binding is displayed as a percentage relative to the DMSO control; (B) specific binding of the mixture is normalized with a baseline set at 0 and a maximal response set at 100% and compared to the expected curve calculated using the CA model. The dotted lines represent the 95% CI for the observed response curve. Values represent Means ± SEM. PCB 95 (n=12), PCB 52 (n=6), mixture (n=6).

The CRC of PCB 95 was significantly altered in the presence of PCB 47 (Figure 3, F=14.41, p < 0.0001). The PCB 95 curve was most impacted when run in the presence of 10 µM PCB 47, which causes minimal activity when run alone. This result suggests that minimally active congeners can compete for the binding site on the RyR, especially at higher concentrations. Finally, the CRC of PCB 95 alone was not significantly different from that observed for PCB 95 in the presence of the RyR1 inactive PCB 126 (Figure 4, F=1.68, p = 0.08). When PCB 126 was present at 10 μM, it caused a slight reduction in the maximal response caused by PCB 95 (Figure 4). If the curve containing10 µM of PCB 126 was removed from the analysis, there was more confidence that the curves were not significantly different (F = 1.27, p = 0.29). As such, the PCB 95 absolute EC_50_ and EC_2X_ did not vary between the curves (Figure 4B and 4C), further suggesting that PCB 126 did not affect PCB 95 RyR1 binding activity.

**Figure 3.**
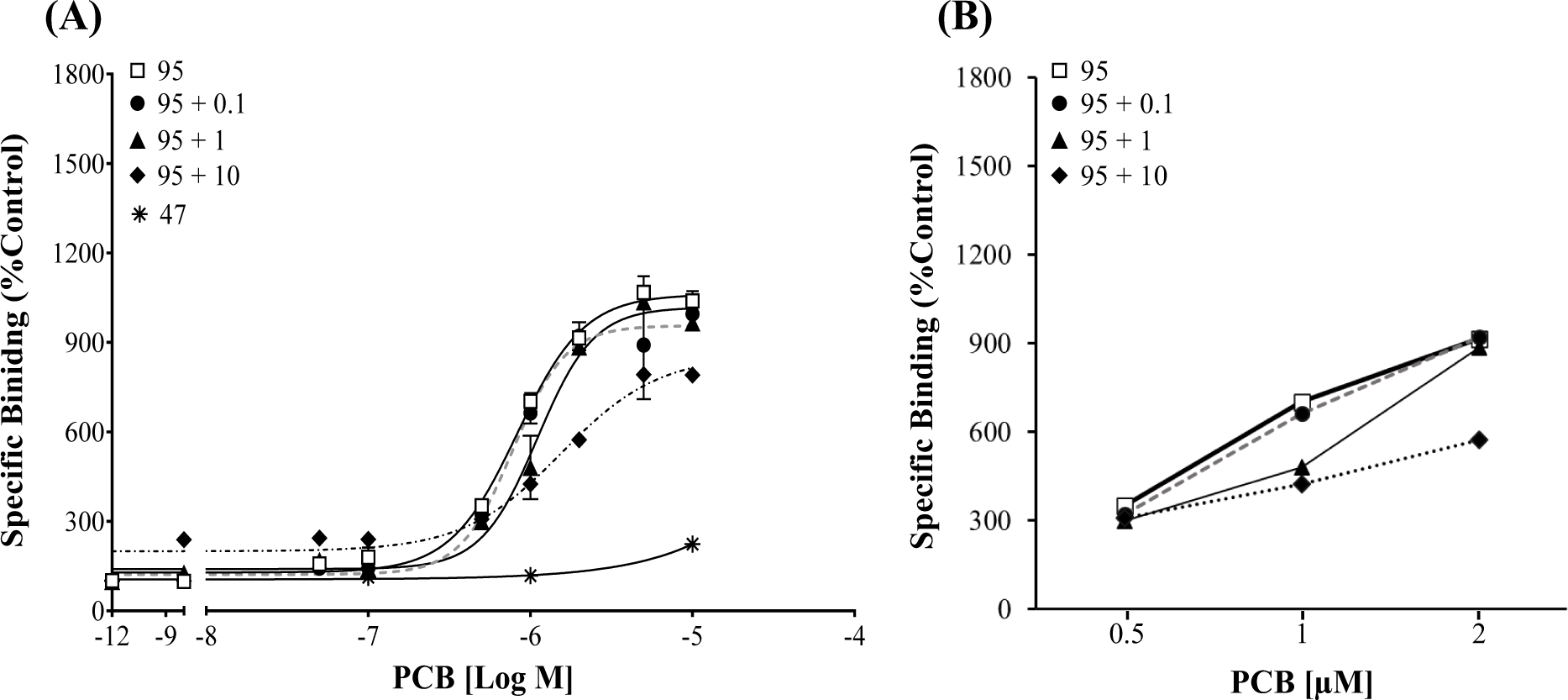
Minimally active PCBs alter the activity of efficacious congener PCB 95. (A) [^3^H]Ry specific binding in the presence of PCB 95 alone or in a mixture with minimally active PCB 47. Binding is displayed as a percentage relative to the DMSO control. (B) Specific binding shown with non-logged concentrations to demonstrate departure from the PCB 95 response curve. Values represent Means ± SEM. PCB 95 (n=12) and PCB 95 +126 (n=3) per concentration.

**Figure 4.**
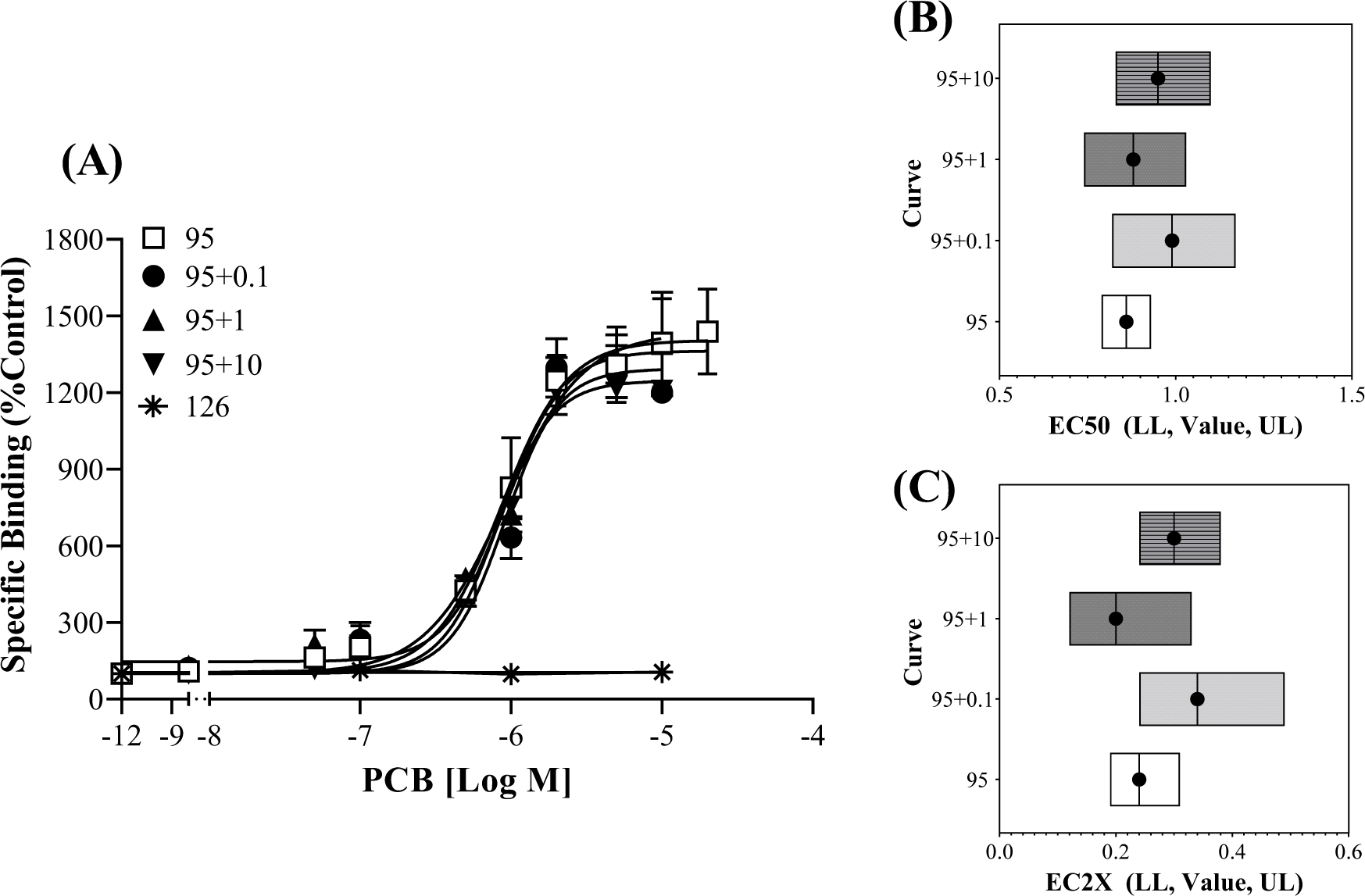
The efficacy and potency of PCB 95 are not altered by the RyR1 inactive congener PCB 126. (A) [^3^H]Ry specific binding in the presence of PCB 95 alone or in a mixture with varying concentrations of PCB 126. Binding is displayed as a percentage relative to the DMSO control and values represent Means ± SEM. PCB 95 (n=12) and PCB 95 +126 (n=3) per concentration. (B) Absolute EC_50_ or (C) EC_2x_ of PCB 95 at the RyR1 when ran alone or with varying concentrations of PCB 126. EC Value with the Lower Limit (LL) and Upper Limit (UL) of the 95%Confidence Interval.

The tertiary mixture containing PCB 47, 51, and 68 (Table S1) activated the RyR, but the maximal response observed was 350% (Figure 5). These findings suggest that congeners PCB 47 and PCB 68, both displaying minimal RyR1 activity, affect the observed activity of the partial agonist PCB 51. This observation is in line with that observed for the activity of PCB 95 in the presence of PCB 47 (Figure 3), suggesting that minimally active congeners compete for the RyR1 binding site. A full CA model expected curve could not be developed for the tertiary mixture due to the minimal activity of PCB 47 and PCB 68. If the CA model was used to develop an EC_2X_ for the tertiary mixture, the expected value was similar but fell outside of the 95% CI of the EC_2X_ observed in the binding assay (Exp: EC_2X_ = 3.34 μM; Obs: EC_2X_ = 2.54 μM, 95% CI 2.27-2.84 μM).

**Figure 5.**
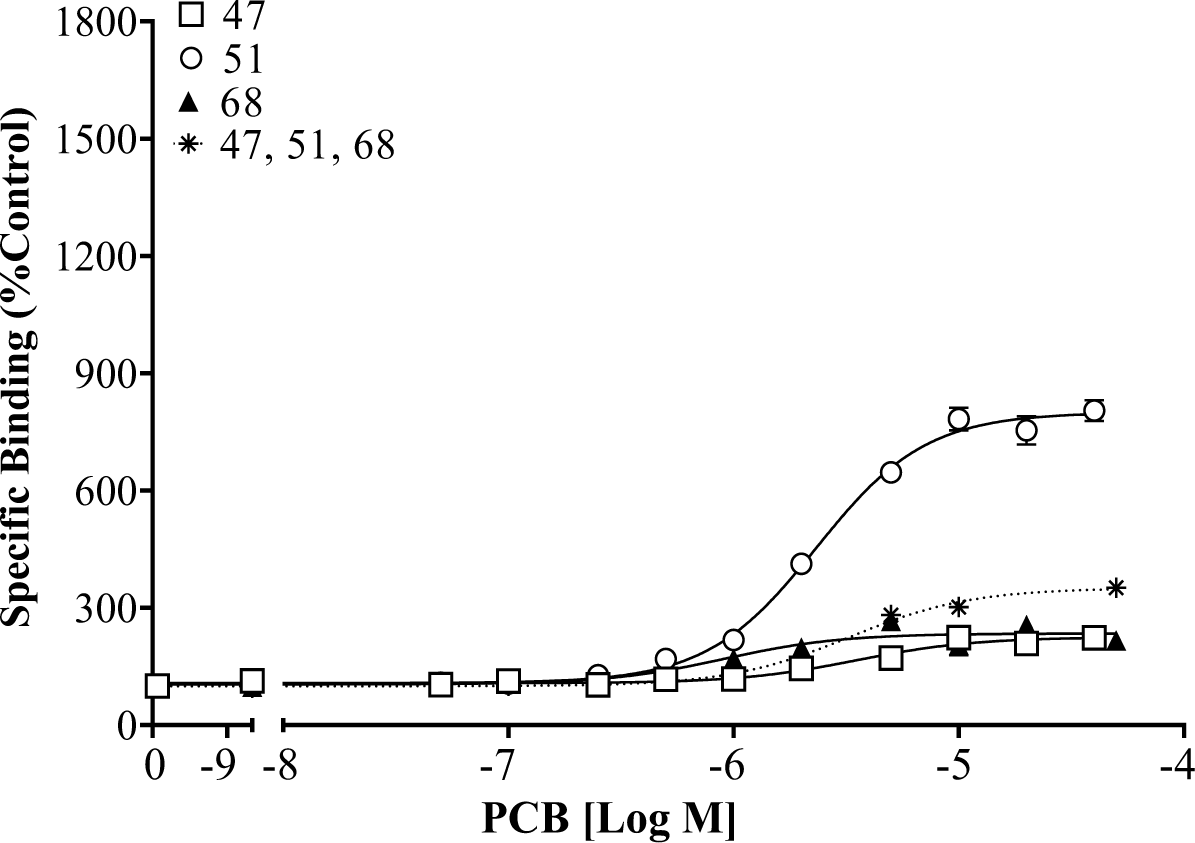
PCB 47, 51 and 68 activate the RyR1 and compete for binding when run in a tertiary mixture. ^3^H]Ry specific binding of PCB 47, 51, and 68 alone or in a tertiary mixture. Binding is displayed as a percentage relative to the DMSO control and values represent Means ± SEM (n=3) for each congener.

### Environmental Mixtures

All the mixtures created to mimic PCBs found in environmental samples or the serum of mothers and children (Table S2-S9) led to significant RyR1 activity (Figure 6). Each mixture had similar maximal effects on the RyR1, causing 400-600% overactivation compared to the DMSO control (Table 2). None of the mixtures caused more than a 700% overactivation of the channel, which would be the activity needed to calculate an absolute EC_50_ normalized to the full agonist PCB 95 (Max 1405%, current study). Therefore, potency was assessed using the EC_2X_ value, where it was found that the mixture detected in the air from schools in the rural area of CJ was the most potent (Table 2). This finding may be due to the presence of the potent congener PCB 52, which represented 17% of the total PCBs found in the CJ schools (Table S3). The level of PCB 52 was also elevated in the EC school air mixture (Table S2, 13%) but was limited in fish (Table S4 and S5) and serum (Table S6-S9). The CA model predicted EC_2X_ value did not overlap with the 95% CI of the observed EC_2X_ of the PCB mixtures CRCs (Table 2). However, the predicted EC_2X_ values were within one magnitude of changing concentration for the observed values.

**Figure 6.**
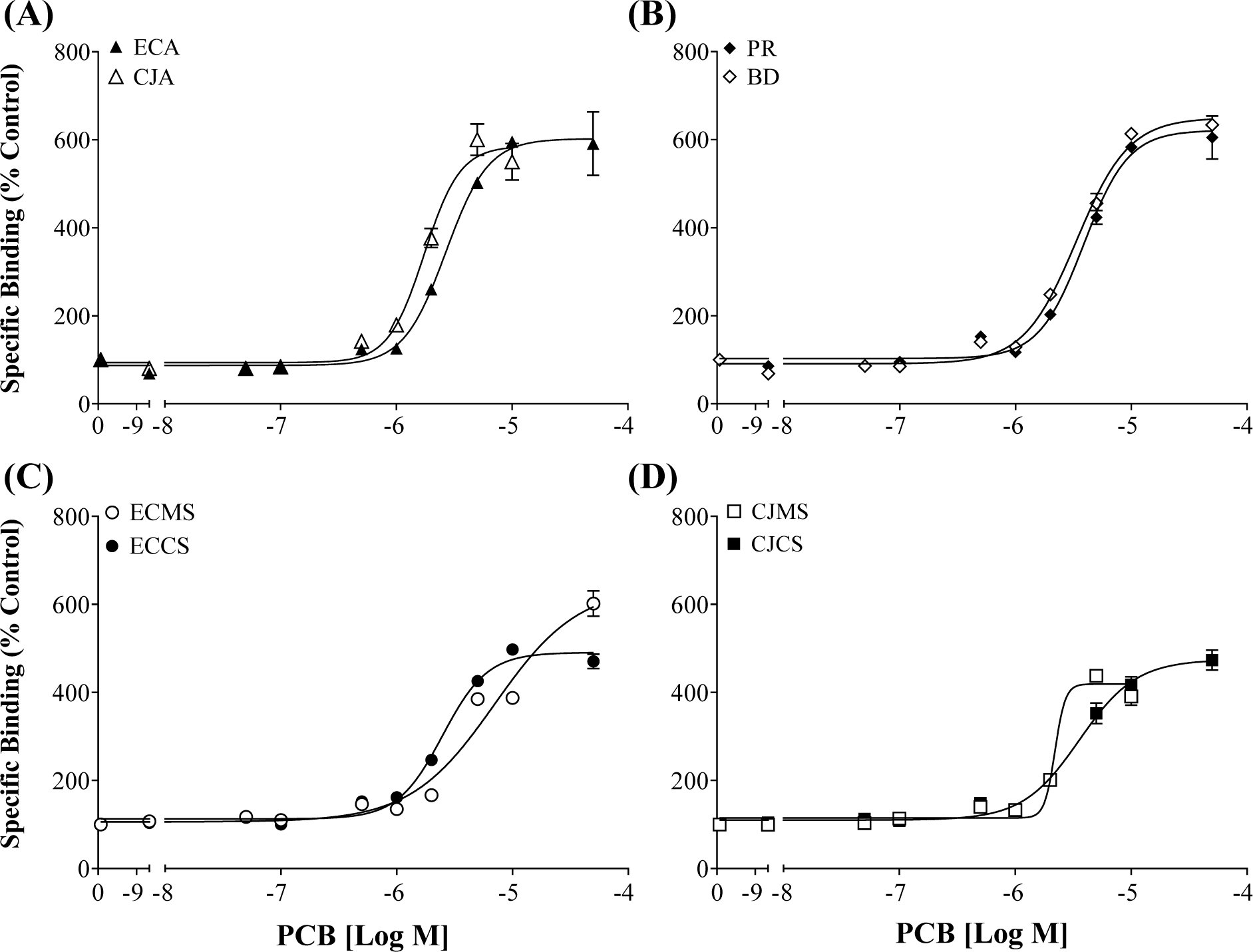
PCBs mixtures found the environment and human serum alter the activity of the RyR1. Binding observed for PCB mixtures mimicking that detected in **(A)** air from schools in East Chicago,(ECA) IN, and Columbus Junction (CJA), IA, **(B)** tissue of predatory (PR) or bottom-dwelling (CD)fish from lakes in Cook County, IL, **(C)** serum from mothers and children from EC (ECMS or ECCS) and **(D)** serum from mothers and children from CJ (CJMS or CJCS). [^3^H] Ry-specific binding is displayed as a percentage relative to the DMSO control. Values represent Means ± SEM, n=3.

**Table 2.** Potency and Efficacy of Environmental or Serum Based PCB Mixtures Toward the Ryanodine Receptor, Isoform 1, in Rabbit Skeletal Muscle. Values in parenthesis represent the 95% Confidence Interval for a given parameter.

| Type of Sample | Details | Max (% Control) | EC<sub>2X</sub> (μM) | CA Model Predicted EC<sub>2X</sub> (μM) |
| --- | --- | --- | --- | --- |
| School Air | East Chicago | 602 (563-642) | 1.56 (1.30-1.85) | 0.61 |
|  | Columbus Junction | 585 (547-631) | 1.10 (0.90-1.30) | 0.52 |
| Lake Fish | Predator | 621 (586-658) | 2.02 (1.72-2.36) | 0.57 |
|  | Bottom Dwelling | 649 (623-678) | 1.60 (1.41-1.82) | 0.59 |
| Serum | East Chicago Mother | 640 (571-781) | 1.93 (1.41-2.53) | 1.10 |
|  | East Chicago Child | 490 (472-510) | 1.45 (1.28-1.63) | 1.13 |
|  | Columbus Junction Mother | 419 (399-440) | 2.00 (1.18-3.78) | 1.09 |
|  | Columbus Junction Child | 474 (447-506) | 1.91 (1.59-2.25) | 1.19 |

PCB mixture concentrations currently detected in the tissue of bottom-dwelling lake fish and the serum of mothers from the urban area of East Chicago cause a significant change in RyR1 activity (Figure 7). All other mixtures would likely not cause significant RyR1 activity when assessed in the ligand binding assay at the concentration detected in the published work (Figure 7). To further assess the applicability of neurotoxic predictive models, observed mixture activity was compared to that predicted for rNEQ bioassay equivalent concentrations (Holland and Pessah, 2021). The rNEQs developed, representing PCB 202 equivalent concentrations, suggested that each mixture assessed in the current study would display RyR activity (Figure 7).

**Figure 7.**
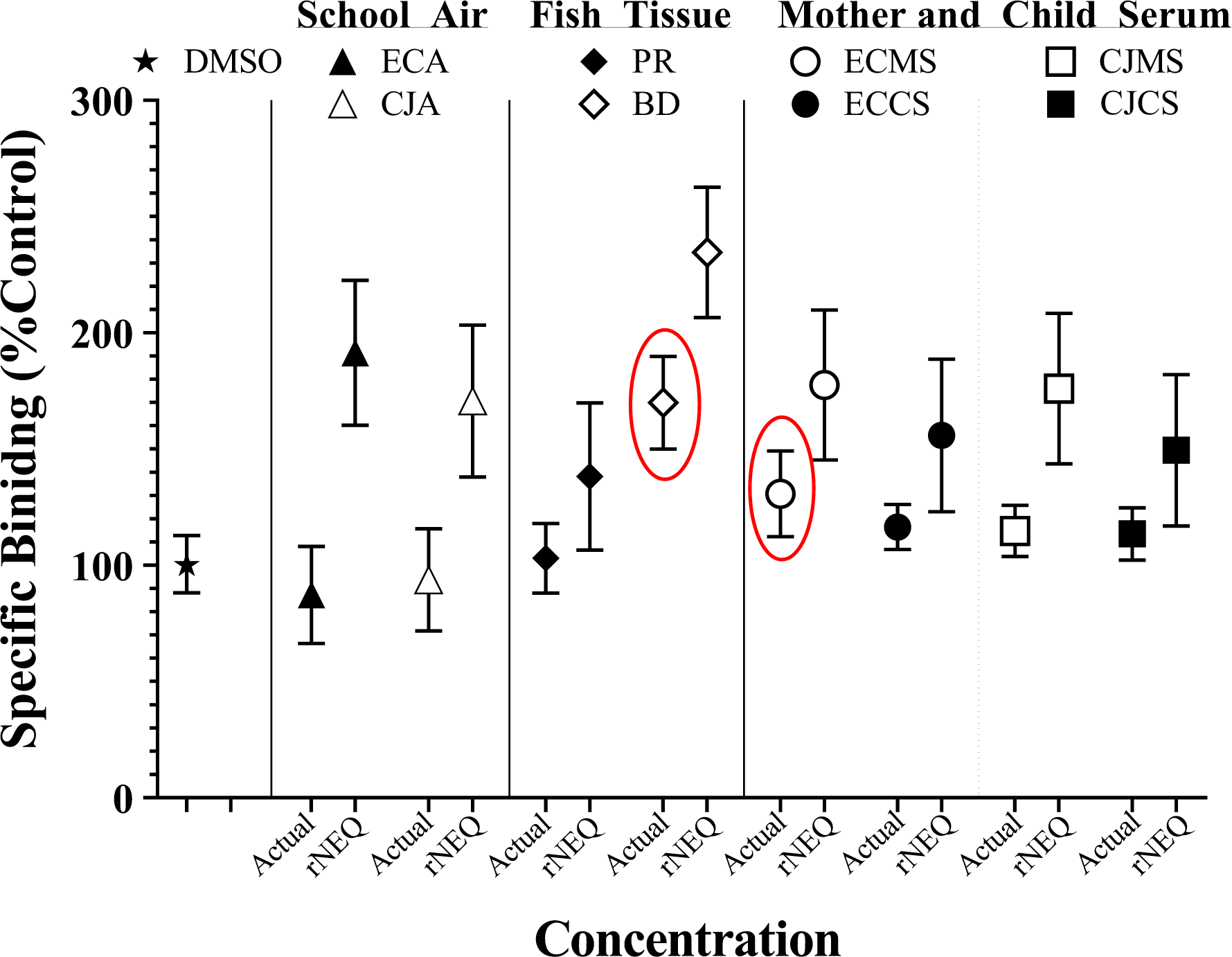
PCB mixtures at concentrations already detected in the environment or human serum alter the activity of the RyR1. Circled symbols represent binding that fell outside the 95% CI of the DMSO control. Actual concentrations are the concentrations of PCB mixtures detected in published work (Table S2-S9). rNEQ concentrations are PCB 202 equivalent concentrations developed with a RyR1 specific NEQ model. Abbreviations: ECA, East Chicago school air; CJA, Columbus Junction school air; Predator, predatory fish tissue; BD, bottom-dwelling fish tissue; ECMS, serum from East Chicago mothers; ECCS, serum from East Chicago children; CJMS, serum from Columbus Junction mothers; CJCS, serum from Columbus Junction mothers.

## DISCUSSION

The current study assessed the activity of NDL PCB congeners as single compounds and when present in binary, tertiary, and complex mixtures. Results on single congeners are in line with previous work (Pessah *et al*., 2006; Holland, *et al*., 2017a). Data show that efficacious congeners display additivity at the mammalian RyR and that minimally active congeners compete with highly potent and efficacious congeners for the RyR binding site. Additionally, all complex mixtures investigated in this study caused RyR overactivation in ligand-binding assays. Neither the CA nor rNEQ models perfectly predicted the potency or RyR1 activity of the complex mixtures, but they were often within a magnitude of change. Importantly, published concentrations of PCB mixtures found in serum caused RyR1 overactivation, highlighting the need to develop an appropriate platform able to predict the neurotoxic risk presented by complex NDL PCB mixtures.

Findings that NDL PCB binary mixtures cause additivity at the mammalian RyR are consistent with work in rainbow trout (Fritsch and Pessah, 2013). However, work in the rainbow trout found that a complex mixture containing 20 PCBs followed the additivity predictions of the CA model, which contrasts with the current study. This difference could be because the rainbow trout work had full CRCs for each of the 20 congeners used in the CA model (Fritsch and Pessah, 2013), whereas the current study needed to use congener EC_2X_ due to a lack of CRC data for all the congeners in each of the published mixtures (Table S1-S9). Additionally, in the trout study, all congeners displayed similar efficacy in binding assays making the data more applicable to the CA model. Finally, teleost fish have additional RyR isoforms compared to mammalian species (Holland, *et al*. 2017b), creating potential differences in binding observed between species. Regardless, both studies support PCB congener additivity at the RyR, especially at low environmentally relevant concentrations.

The CA model predicted environmental mixtures would be more potent (i.e., lower EC values) than those observed with mixture CRCs. There are several limitations to the use of the CA model, which assume that all chemicals in a mixture display similar efficacy and produce parallel response curves (Howard et al., 2010; Howard and Webster, 2009; Silva et al., 2002). The generalized concentration addition (GCA) model is an extension of the CA model, which can be applied to mixtures with full and partial agonists and antagonists (Howard and Webster, 2009; Howard *et al*., 2010). The GCA requires a concentration-response function for each chemical in a mixture (de la Rosa et al., 2020; Howard et al., 2010), which is a challenge for NDL PCBs given the number known or predicted to display RyR activity (Holland et al., 2017a; Pessah et al., 2006; Rayne and Forest, 2010). Work using GCA, to date, often uses extensive combinations of two compounds of interest and has not compared GCA predictions to observed effects of complex environmentally relevant mixtures (de la Rosa et al., 2020; Howard et al., 2010). Additional work has proposed using the CA model in conjunction with the Independent Action model (CA/IA model) to assess complex mixture effects (Escher et al., 2020, 2018). Here, the proposed CA/IA model uses low effect levels on non-log transformed CRCs to predict mixture effects. These predictions would be more relevant for the lower concentrations of chemicals detected in the environment and include chemicals acting on a diverse set of molecular targets (Escher et al., 2020). Applying further mixture models to predict NDL PCB-induced neurotoxicity would greatly contribute to hazard assessment development. Further, work should include other RyR active chemicals, such as polybrominated diphenyl ether (Kim et al., 2011), that likely display additivity with NDL PCBs at the RyR (Fritsch and Pessah, 2013).

NDL PCBs comprise more than 60% of the total PCB concentration in environmental mixtures (Table S2-S9). In this study, PCB mixtures were prepared to include the 25 PCB congeners with the highest concentration detected in the environment or human serum samples, where many of the congeners contributing the largest percentage of total PCBs displaying RyR1 activity (e.g., PCB 52 or PCB 180). Importantly, mixtures utilized in the current study cause RyR activity at concentrations detected in the environment or human serum. This is of particular concern for PCB profiles found in the serum of mothers and children (Marek et al. 2013), where such exposure has been associated with developmental neurotoxicity (reviewed by Pessah et al. 2019). Currently, there is limited data regarding levels of PCBs in brain tissue, but they do show that highly chlorinated congeners were predominant, including RyR active PCB 153 and 180. However, most studies measure a limited number of congeners (Chu et al., 2003; Dewailly et al., 1999; Mitchell et al., 2012) and often work with adult tissue, hindering an understanding of brain PCB profiles and potential impacts across age groups. A more recent study, measuring all PCBs and OH-PCBs, found that different congeners predominate in different brain regions of postmortem patients and that these patterns vary widely between age classes (Li et al. 2022). Here, adult patients tended to have more consistent patterns between brain regions, which all contained highly chlorinated congeners. Conversely, neonatal brains had variable patterns across regions, and the tissue had a higher percentage of lower chlorinated PCBs. The profile in the human brain likely differentially contributes to neurotoxicity but more PCB data in brain tissue should include extensive detection of congeners and their metabolites.

Considerable research has connected NDL PCB activity toward the RyR and other potential molecular targets, such as thyroid hormone-dependent mechanisms, neurotransmitter release, or activity at the GABA receptor (reviewed by Pessah et al. 2019; Klocke and Lein 2020; Klocke et al. 2020) with neurotoxicity. Each congener likely displays varying toxicity toward different molecular targets (Holland and Pessah, 2021; Klocke et al. 2020), but activity through varying targets may converge to cause similar cellular impacts (Klocke et al. 2020). Here, neurotoxic outcomes have been noted for both legacy PCBs, which were part of Aroclor formulations, and the non-Aroclor contemporary congeners that are byproducts of pigment manufacturing (Jahnke and Hornbuckle, 2019) or finishing of commercial products such as wood cabinets (Herkert et al., 2018). Notably, the non-Aroclor PCB 11, detected in outdoor and indoor air, and the plasma of pregnant women (Sethi et al., 2017) does not display RyR activity (Holland et al. 2017a). However, PCB 11 alters dendritic growth through CREB-dependent mechanisms, and this effect is not due to impacts on cellular Ca^2+^ levels (Sethi et al., 2018), as seen with higher chlorinated PCBs. Specifically, RyR1 active PCB 95 alters CREB-mediated gene expression in a Ca^2+^-dependent manner, contributing to altered dendritic growth (Wayman et al., 2012). Work has yet to directly address the cellular or organismal effect of PCB95 in a mixture with PCB 11, but together they likely cause distinct changes in CREB-mediated development. As such, predictive models must consider multiple molecular targets to adequately predict the neurotoxic impacts of NDL PCBs and structurally related compounds (Holland and Pessah, 2021; Pradeep et al., 2019). And while there is a need for toxicity assessments on individual PCB congeners, more work should test complex mixtures of NDL PCBs and related compounds relevant to environmental and human samples (Klocke et al., 2020).

## CRediT authorship contribution statement

**Justin Griffin:** Methodology, Investigation, Validation, Formal Analysis, Writing-original draft, Funding acquisition. **Xueshu Li:** Methodology, Investigation, Formal Analysis, Writing-original draft, Writing-review &editing. **Hans-Joachim Lehmler:** Conceptualization, Project Administration, Supervision, Funding acquisition, Resources, Writing-original draft, Writing-review &editing. **Erika Holland:** Conceptualization, Methodology, Validation, Data curation, Visualization, Project Administration, Supervision, Funding acquisition, Resources, Writing-original draft, Writing-review &editing.

## Declaration of Generative AI and AI-assisted technologies in the writing process

Material presented in this work was created by the authors. Writing and related figures and tables were not generated by AI and AI was not used to assist in the creation of this manuscript.

## Supporting information

PCB Mixture Supplement

## Acknowledgments

Appreciation is extended to Dr. Isaac N. Pessah and Dr. Wei Feng from UC Davis for the generous JSR gift, Dr. Leanne Stahl from the US Environmental Protection Agency (Office of Water; Washington, DC) for supplying the raw PCB concentrations found in the tissue of fish from US lakes, and Dr. Ram Dhakal from University of Iowa for the formulation of the synthetic PCB mixtures.

## Supplementary File

Additional materials related to preparing and authenticating PCB mixtures relevant to environmental and human samples can be found in the supplemental file available online.

