## Supplementary material for "Predicted Versus Observed Activity of PCB Mixtures Toward the Ryanodine Receptor": PCB Mixture Supplement

**Table of Contents**

*Sources of individual PCB congeners* ***S3***

*General procedure for the preparation of environmental PCB mixtures* ***S3***

*Authentication of PCB mixtures* ***S4***

***Table S1**. Percentage contribution of PCB 47, 51, and 68 to the total PCBs found in the indoor air of new construction buildings.* ***S5***

***Table S2.** PCB congeners found in the air of schools in the urban area of east Chicago, IN* ***S6***

***Table S3**. PCB congeners found in the air of schools from Columbus Junction, IA* ***S7***

***Table S4**. PCB congeners found in the tissue of predatory lake fish from Cook County, IL* ***S8***

***Table S5.** PCB congeners found in the tissue of bottom dwelling lake fish from Cook County, IL* ***S9***

***Table S6**. PCB congeners found in the serum of mothers from the urban area of east Chicago, IN* ***S10***

***Table S7***. *PCB congeners found in the serum of children from the urban area of east Chicago, IN* **S11**

***Table S8.*** *PCB congeners found in the serum of mothers from the rural area of Columbus Junction, IA* **S12**

***Table S9.*** *PCB congeners found in the serum of children from the rural area of Columbus Junction, IA* **S13**

***Table S10.*** *Similarity coefficient (cos Θ) between the profiles of targeted PCB mixture and the lab synthesized mimic mixture.* **S14**

*References* **S15**

### Sources of individual PCB congeners

Pesticide-grade solvents were used to prepare the desired solution. PCB 1, PCB 3, PCB 20, PCB 28, PCB 47, PCB 52, PCB 84, PCB 101, PCB 118, PCB 138, PCB 149, PCB 153, and PCB 180 were synthesized and characterized as described previously (Kania-Korwel et al., 2004; Lehmler et al., 2005; Lehmler and Robertson, 2001; Sethi et al., 2019; Telu et al., 2010; Vyas et al., 2006). PCB 4 (lot: 990505LB-AC), PCB 8 (lot: 9810011B-AC-01), PCB 18 (lot: 09300110KS), PCB 21 (lot: 081799C-AC-01), PCB 31(lot: 27202), PCB 33 (lot: 012693-01), PCB 37 (lot: 11230-01) , PCB 40 (lot: 920724), PCB 44 (lot: 070899MT-AC), PCB 49 (lot: 190594-01), PCB 51 (lot: 980511LB-AC), PCB 64 (lot:97063LB-AC-01), PCB 66 (lot: 090913DT), PCB 68 (lot:18498) PCB 70 (lot: 20770), PCB 71 (lot: 051809KS-2), PCB 83 (lot: 102809KS, 12260-01), PCB 85 (lot: 931020-01), PCB 87 (lot: 18568), PCB 91 (lot: 980526LB-AC-01), PCB 92 (lot: 010294A), PCB 95 (lot: 100300MT-AC-01), PCB 97 (lot: 22981), PCB 99 (lot: 1270), PCB 105 (lot: 112712KC), PCB 110 (lot: 27084, 27084), PCB 132 (lot: 092613DT), PCB 141 (lot: 70000), PCB 146 (lot: 040111KS), PCB 151 (lot: 052708RC1-01), PCB 156 (lot: 0613131AG), PCB 163 (lot: 01041-01), PCB 170 (lot: 980708LB-AC), PCB 174 (lot: 121004MT-AC), PCB 177 (lot: 951103LPB-AC-01), PCB 183 (lot: 960607LB-AC), PCB 187 (lot: 28524, 20726), PCB 194 (lot: 951208LPB), PCB 199 (lot: 940719), PCB 203 (lot: 052593), and PCB 206 (lot: 032713KC) were bought from AccuStandard (New Haven, CT, USA).

### General procedure for the preparation of environmental PCB mixtures

Environmental PCB mixtures were prepared by dissolving the desired amount of each PCB congener in acetone in a volumetric flask. The solutions containing individual PCB congeners were mixed in the appropriate ratios to generate the PCB mixtures. Each mixture solution was divided into four different aliquots and the solvent was completely evaporated under a gentle stream of ultrapure nitrogen gas. Each aliquot contained 3 to 5 mg of the neat PCB mixtures.

**Authentication of PCB mixtures**

All synthetic PCB mixtures were authenticated (Li et al., 2018) with a gas chromatography-tandem mass spectrometry (GC-MS/MS) method that allows for the identification and quantification of all 209 PCB congeners as 174 individual or co-eluting chromatographic peaks(Li et al., 2022). Because synthetic PCB congeners can contain small impurities of other PCB congeners which may be the biologically active congeners (Li et al., 2018; Zhang et al., 2022), this approach allows us also to quantify these impurities. Briefly, one aliquot of each PCB mixture was dissolved in 1 mL of hexane. Twenty to 100 µL (depending on the concentration of individual PCB congeners) of the solution were transferred to a 100 mL volumetric flask and diluted with isooctane to yield solutions with PCB concentrations ranging from 5 to 634 ng/mL. One mL of the isooctane solution was spiked with internal standards (250 ng d-PCB 30 and 500 ng PCB 204). The diluted samples were analyzed on a GC-MS/MS system (Agilent 7890A GC system, Agilent 7000 Triple Quad, Agilent 7693 autosampler) on an SPB-Octyl column (30 m length, 250 µm inner diameter, and 0.25 µm film thickness; Sigma-Aldrich, St. Louis, MO, USA) in multiple reaction monitoring mode (MRM). Instrument blanks of isooctane were analyzed before and after calibration and after the samples to ensure no carryover.

Comparisons of the environmental target PCB profiles and the corresponding synthetic PCB profiles are shown in Tables S1 to S9. The experimental PCB congener profiles of the target congeners were compared to the target profiles using the similarity coefficient (cos Θ). The cos Θ is frequently used to compare PCB profiles and ranges from 0 to 1 (Li et al., 2022, 2020). A cos Θ of 1 indicates that the profiles are identical, whereas a cos Θ of 0 indicates that the profiles are completely different. As shown in Table S10, the synthetic PCB mixtures are a good match with the environmental target profiles.

**Table S1.** Percentage contribution of PCB 47, 51, and 68 to the total PCBs found in the indoor air of new construction buildings (Herkert et al., 2018). Congeners in red are impurities present in the neat PCB congeners used to prepare this mixture.

| **(A) Concentrations Detected in Indoor Air** | | | **(B) Synthetic PCB Congener Profile** | |
|---|---|---|---|---|
| **Congener** | **Concentration ($ng/m^3$)** | **Percent of Total** | **Congener** | **Percent of Total** |
| PCB 20+28 | | | PCB 20+28 | 0.05 |
| PCB 21+33 | | | PCB 21+33 | 0.18 |
| PCB 44+47+65 | 1.88 | 70.10 | PCB 44+47+65 | 79.50 |
| PCB 49+69 | | | PCB 49+69 | 0.48 |
| PCB 51 | 0.53 | 19.81 | PCB 51 | 13.57 |
| PCB 68 | 0.27 | 10.09 | PCB 68 | 6.23 |
| **Sum of 47/51/68** | **2.68** | **100.00** | **Sum of 47/51/68** | **100.01** |
| *Sum All Congeners* | *5.03* | | | |
| *Congeners % of Total* | *53.20* | | | |

**Table S2.** PCB congeners found in the air of schools in the urban area of East Chicago, IN (Marek et al., 2017). Congeners in red are impurities present in the neat PCB congeners used to prepare this mixture.

| **(A) Top 25 Congers Detected in Air** | | | **(B) Synthetic PCB Congener Profile** | |
|---|---|---|---|---|
| **Congener** | **Amount (ng/m$^3$)** | **Percent of Total** | **Congener** | **Percent of Total** |
| PCB 1 | 7.23 | 4.77 | PCB 1 | 3.86 |
| PCB 3 | 5.05 | 3.33 | PCB 3 | 2.39 |
| PCB 4 | 3.30 | 2.18 | PCB 4 | 2.04 |
| PCB 8 | 7.28 | 4.81 | PCB 8 | 4.39 |
| PCB 18+30 | 2.92 | 1.93 | PCB 18+30 | 1.40 |
| PCB 20+28 | 2.81 | 1.86 | PCB 20+28 | 1.47 |
| PCB 26+29 | | | PCB 26+29 | 0.02 |
| PCB 31 | 2.89 | 1.91 | PCB 31 | 1.75 |
| PCB 37 | | | PCB 37 | 0.04 |
| PCB 44+47+65 | 6.76 | 4.46 | PCB 44+47+65 | 4.19 |
| PCB 49+69 | 4.67 | 3.09 | PCB 49+69 | 3.05 |
| PCB 52 | 19.78 | 13.06 | PCB 52 | 11.48 |
| PCB 56 | | | PCB 56 | 0.04 |
| PCB 61+70+ 74+76 | 10.46 | 6.91 | PCB 61+70+ 74+76 | 6.03 |
| PCB 64 | 2.11 | 1.40 | PCB 64 | 3.67 |
| PCB 66 | 2.39 | 1.58 | PCB 66 | 1.26 |
| PCB 84 | 3.97 | 2.62 | PCB 84 | 2.42 |
| PCB 86+97+109+119 | 2.92 | 1.93 | PCB 86+97+109+119 | 1.96 |
| PCB 87+125 | 4.98 | 3.29 | PCB 87+125 | 3.30 |
| PCB 90+101+113 | 13.20 | 8.72 | PCB 90+101+113 | 8.34 |
| PCB 92 | 2.33 | 1.54 | PCB 92 | 6.84 |
| PCB 95 | 13.62 | 9.00 | PCB 95 | 8.64 |
| PCB 99 | 4.91 | 3.24 | PCB 99 | 3.44 |
| PCB 110 | 10.83 | 7.15 | PCB 110 | 7.44 |
| PCB 118 | 5.89 | 3.89 | PCB 118 | 4.06 |
| PCB 129+138+160 | 3.83 | 2.53 | PCB 129+138+160 | 2.10 |
| PCB 135+ 151 | | | PCB 135+ 151 | 0.06 |
| PCB 146 | | | PCB 146 | 0.03 |
| PCB 147+ 149 | 4.36 | 2.88 | PCB 147+ 149 | 2.59 |
| PCB 153+168 | 2.93 | 1.93 | PCB 153+168 | 1.65 |
| PCB 174 | | | PCB 174 | 0.02 |
| PCB 182 | | | PCB 182 | 0.02 |
| PCB 184 | | | PCB 184 | 0.02 |
| PCB 188 | | | PCB 188 | 0.02 |
| **ΣPCB** | **151.41** | **100.00** | **ΣPCB** | **100.03** |
| *Sum of 209 Congeners* | *193.42* | | | |
| *Congener % of Total* | *78.28* | | | |

**Table S3.** PCB congeners found in the air of schools from Columbus Junction, IA (Marek et al., 2017). Congeners in red are impurities present in the neat PCB congeners used to prepare this mixture.

| **(A) Top 25 Congers Detected in Air** | | | **(B) Synthetic PCB Congener Profile** | |
|---|---|---|---|---|
| **Congener** | **Amount (ng/m$^3$)** | **Percent of Total** | **Congener** | **Percent of Total** |
| PCB 18+30 | 1.29 | 1.78 | PCB 18+30 | 1.44 |
| PCB 20+28 | 1.12 | 1.55 | PCB 20+28 | 1.27 |
| PCB 26+29 | | | PCB 26+29 | 0.01 |
| PCB 31 | 1.49 | 2.06 | PCB 31 | 1.89 |
| PCB 40+71 | 0.75 | 1.04 | PCB 40+71 | 1.16 |
| PCB 44+47+65 | 4.07 | 5.63 | PCB 44+47+65 | 5.63 |
| PCB 49+69 | 1.99 | 2.75 | PCB 49+69 | 2.87 |
| PCB 52 | 12.06 | 16.67 | PCB 52 | 18.00 |
| PCB 61+70+74+76 | 5.84 | 8.08 | PCB 61+70+74+76 | 8.52 |
| PCB 64 | 1.41 | 1.95 | PCB 64 | 5.19 |
| PCB 66 | 1.42 | 1.97 | PCB 66 | 1.79 |
| PCB 81 | | | PCB 81 | 0.04 |
| PCB 84 | 1.95 | 2.69 | PCB 84 | 2.62 |
| PCB 86+98+109+119 | 1.56 | 2.15 | PCB 86+98+109+119 | 2.13 |
| PCB 87+125 | 2.27 | 3.14 | PCB 87+125 | 2.80 |
| PCB 90+101+113 | 7.77 | 10.75 | PCB 90+101+113 | 9.45 |
| PCB 91 | 0.97 | 1.33 | PCB 91 | 1.78 |
| PCB 92 | 1.34 | 1.85 | PCB 92 | 1.82 |
| PCB 95 | 7.39 | 10.21 | PCB 95 | 10.60 |
| PCB 99 | 2.88 | 3.98 | PCB 99 | 3.56 |
| PCB 110 | 5.27 | 7.29 | PCB 110 | 6.30 |
| PCB 118 | 2.63 | 3.64 | PCB 118 | 3.33 |
| PCB 129+138+160 | 1.55 | 2.14 | PCB 129+138+160 | 1.52 |
| PCB 132 | 0.76 | 1.05 | PCB 132 | 0.85 |
| PCB 135+151 | 0.88 | 1.22 | PCB 135+151 | 1.04 |
| PCB 146 | | | PCB 146 | 0.02 |
| PCB 147+149 | 2.19 | 3.03 | PCB 147+149 | 2.38 |
| PCB 153+168 | 1.47 | 2.03 | PCB 153+168 | 1.69 |
| PCB 174 | | | PCB 174 | 0.03 |
| PCB 181 | | | PCB 181 | 0.03 |
| PCB 182 | | | PCB 182 | 0.07 |
| PCB 184 | | | PCB 184 | 0.09 |
| PCB 186 | | | PCB 186 | 0.01 |
| PCB 188 | | | PCB 188 | 0.05 |
| PCB 209 | | | PCB 209 | 0.02 |
| **ΣPCB** | **72.31** | **100.00** | **ΣPCB** | **100.00** |
| *Sum of 209 Congeners* | *82.94* | | | |
| *Congener % of Total* | *83.17* | | | |

**Table S4.** PCB congeners found in the tissue of predatory lake fish from Cook County, IL (Stahl et al., 2009). Congeners in red are impurities present in the neat PCB congeners used to prepare this mixture.

| **(A) Top 25 Congers Detected in Air** | | | **(B) Synthetic PCB Congener Profile** | |
|---|---|---|---|---|
| **Congener** | **Amount ($ng/m^3$)** | **Percent of Total** | **Congener** | **Percent of Total** |
| PCB 20+28 | 517 | 1.70 | PCB 20+28 | 1.69 |
| PCB 49+69 | 476 | 1.57 | PCB 49+69 | 2.11 |
| PCB 52 | 803 | 2.64 | PCB 52 | 3.24 |
| PCB 61+70+74+76 | 655 | 2.16 | PCB 61+70+74+76 | 2.29 |
| PCB 67 | | | PCB 67 | 0.01 |
| PCB 79 | | | PCB 79 | 0.03 |
| PCB 83+99 | 998 | 3.28 | PCB 83 | 1.36 |
| | | | PCB 99 | 2.04 |
| PCB 86+87+97+108+119+125 | 773 | 2.54 | PCB 86+87+97+109+119 | 2.97 |
| PCB 90+101+112 | 1650 | 5.43 | PCB 90+101+113 | 5.09 |
| PCB 93+95+98+100+102 | 507 | 1.67 | PCB 95 | 1.75 |
| PCB 105 | 476 | 1.57 | PCB 105 | 1.17 |
| PCB 110+115 | 1170 | 3.85 | PCB 110+115 | 3.12 |
| PCB 118 | 1290 | 4.24 | PCB 118 | 4.61 |
| PCB 128+166 | | | PCB 128+166 | 0.10 |
| PCB 129+138+163 | | | PCB 129+138+163 | 0.07 |
| PCB 130 | | | PCB 130 | 0.04 |
| PCB 135+151+154 | 702 | 2.31 | PCB 135+151 | 2.11 |
| PCB 141 | 670 | 2.20 | PCB 141 | 2.24 |
| PCB 146 | 1010 | 3.32 | PCB 146 | 2.62 |
| PCB 147+149 | 1930 | 6.35 | PCB 147+149 | 5.49 |
| PCB 153+168 | 4560 | 15.01 | PCB 153+168 | 14.94 |
| PCB 167 | | | PCB 167 | 0.03 |
| PCB 170 | 1230 | 4.05 | PCB 170 | 3.81 |
| PCB 172 | | | PCB 172 | 0.05 |
| PCB 174 | 607 | 2.00 | PCB 174 | 1.67 |
| PCB 177 | 535 | 1.76 | PCB 177 | 4.50 |
| PCB 178 | | | PCB 178 | 0.02 |
| PCB 180+193 | 3620 | 11.91 | PCB 180+193 | 10.69 |
| PCB 181 | | | PCB 181 | 0.03 |
| PCB 182 | | | PCB 182 | 0.08 |
| PCB 183+185 | 1180 | 3.88 | PCB 183 | 3.36 |
| PCB 184 | | | PCB 184 | 0.08 |
| PCB 187 | 2670 | 8.79 | PCB 187 | 9.24 |
| PCB 188 | | | PCB 188 | 0.05 |
| PCB 189 | | | PCB 189 | 0.02 |
| PCB 194 | 625 | 2.06 | PCB 194 | 2.07 |
| PCB 195 | | | PCB 195 | 0.01 |
| PCB 198 | 1060 | 3.49 | PCB 198+199 | 3.23 |
| PCB 202 | | | PCB 202 | 0.01 |
| PCB 203 | 675 | 2.22 | PCB 203 | 1.93 |
| PCB 296 | | | PCB 296 | 0.01 |
| PCB 209 | | | PCB 209 | 0.02 |
| **ΣPCB** | **30389** | **100.00** | **ΣPCB** | **100.00** |
| *Sum of 209 Congeners* | *42591* | | | |
| *Congeners % of Total* | *71.35* | | | |

**Table S5.** PCB congeners found in the tissue of bottom dwelling lake fish from Cook County, IL (Stahl et al., 2009). Congeners in red are impurities present in the neat PCB congeners used to prepare this mixture.

| (A) Top 25 Congers Detected in Fish Tissue | | | (B) Synthetic PCB Congener Profile | |
|---|---|---|---|---|
| **Congener** | **Amount (ng/kg)** | **Percent of Total** | **Congener** | **Percent of Total** |
| PCB 49+69 | | | PCB 49+69 | 0.01 |
| PCB 50+53 | | | PCB 50+53 | 0.01 |
| PCB 52 | 6010 | 1.25 | PCB 52 | 1.43 |
| PCB 61+70+74+ 76 | 8620 | 1.79 | PCB 61+70+74+ 76 | 1.90 |
| PCB 79 | | | PCB 79 | 0.02 |
| PCB 83+99 | 12700 | 2.64 | PCB 83 | 1.22 |
| | | | PCB 99 | 1.07 |
| PCB 86+87+97+108+119+125 | 10600 | 2.20 | PCB 86+87+97+108+119+125 | 2.45 |
| PCB 90+101+113 | 21500 | 4.47 | PCB 90+101+113 | 3.84 |
| PCB 93+95+98+100+102 | 7140 | 1.48 | PCB 93+95+98+100+102 | 1.54 |
| PCB 110+115 | 13900 | 2.89 | PCB 110+115 | 2.29 |
| PCB 118 | 15300 | 3.18 | PCB 118 | 3.58 |
| PCB 129+138+160+163 | 58200 | 12.10 | PCB 129+138+160+163 | 12.70 |
| PCB 132 | 7740 | 1.61 | PCB 132 | 1.67 |
| PCB 133 | | | PCB 133 | 0.02 |
| PCB 135+151+154 | 9260 | 1.92 | PCB 135+151+154 | 1.74 |
| PCB 141 | 9890 | 2.06 | PCB 141 | 2.20 |
| PCB 146 | 13000 | 2.70 | PCB 146 | 2.01 |
| PCB 147+149 | 29100 | 6.05 | PCB 147+149 | 4.92 |
| PCB 153+168 | 75600 | 15.72 | PCB 153+168 | 15.24 |
| PCB 156+157 | | | PCB 156+157 | 0.03 |
| PCB 159 | | | PCB 159 | 0.02 |
| PCB 167 | | | PCB 167 | 0.03 |
| PCB 169 | | | PCB 169 | 0.31 |
| PCB 170 | 16900 | 3.51 | PCB 170 | 3.31 |
| PCB 172 | | | PCB 172 | 0.05 |
| PCB 174 | 10600 | 2.20 | PCB 174 | 1.60 |
| PCB 177 | 10600 | 2.20 | PCB 177 | 5.74 |
| PCB 180+193 | 53700 | 11.16 | PCB 180+193 | 9.97 |
| PCB 182 | | | PCB 182 | 0.07 |
| PCB 183+185 | 16100 | 3.35 | PCB 183+185 | 2.95 |
| PCB 187 | 31200 | 6.49 | PCB 187 | 6.68 |
| PCB 189 | | | PCB 189 | 0.02 |
| PCB 194 | 11700 | 2.43 | PCB 194 | 2.51 |
| PCB 196 | | | PCB 196 | 0.03 |
| PCB 198+199 | 15500 | 3.22 | PCB 198+199 | 3.26 |
| PCB 203 | 9300 | 1.93 | PCB 203 | 1.73 |
| PCB 206 | 6880 | 1.43 | PCB 206 | 1.81 |
| PCB 209 | | | PCB 209 | 0.02 |
| **ΣPCB** | **481040** | **100.00** | **ΣPCB** | **100.00** |
| *Sum of 209 Congeners* | *6068632* | | | |
| *Congeners % of Total* | *79.27* | | | |

**Table S6.** PCB congeners found in the serum of mothers from the urban area of east Chicago, IN (Marek et al., 2013). Congeners in red are impurities present in the neat PCB congeners used to prepare this mixture.

| (A) Congers Detected above the LOQ in Serum | | | (B) Synthetic PCB Congener Profile | |
|---|---|---|---|---|
| **Congener** | **Amount (ng/g lipid)** | **Percent of Total** | **Congener** | **Percent of Total** |
| PCB 8 | 2.44 | 1.25 | PCB 8 | 1.24 |
| PCB 18+30 | 2.27 | 1.16 | PCB 18+30 | 0.72 |
| PCB 20+28 | 5.39 | 2.76 | PCB 20+28 | 1.97 |
| PCB 31 | 3.64 | 1.87 | PCB 31 | 1.93 |
| PCB 49+69 | 2.39 | 1.22 | PCB 49+69 | 0.92 |
| PCB 66 | 5.36 | 2.75 | PCB 66 | 1.86 |
| PCB 77 | | | PCB 77 | 0.05 |
| PCB 83 | 14.85 | 7.61 | PCB 83 | 5.30 |
| PCB 84 | 2.92 | 1.50 | PCB 84 | 1.23 |
| PCB 90+101+ 113 | | | PCB 90+101+ 113 | 0.16 |
| PCB 99 | 4.29 | 2.20 | PCB 99 | 1.62 |
| PCB 105 | 3.52 | 1.80 | PCB 105 | 1.59 |
| PCB 110+115 | 6.92 | 3.55 | PCB 110+115 | 3.87 |
| PCB 118 | 14.47 | 7.41 | PCB 118 | 10.77 |
| PCB 128+166 | | | PCB 128+166 | 0.10 |
| PCB 129+137+138+163+164 | 26.46 | 13.56 | PCB 129+138+163 | 12.66 |
| PCB 146 | 3.51 | 1.80 | PCB 146 | 1.34 |
| PCB 147+149 | | | PCB 147+149 | 0.09 |
| PCB 153+168 | 35.67 | 18.28 | PCB 153+168 | 28.30 |
| PCB 156+157 | 6.52 | 3.34 | PCB 156+157 | 3.51 |
| PCB 169 | | | PCB 169 | 0.15 |
| PCB 170 | 10.41 | 5.33 | PCB 170 | 2.91 |
| PCB 180+193 | 24.38 | 12.49 | PCB 180+193 | 6.96 |
| PCB 183 | 1.93 | 0.99 | PCB 183 | 0.84 |
| PCB 187 | 9.08 | 4.65 | PCB 187 | 7.25 |
| PCB 198+199 | 6.1 | 3.13 | PCB 198+199 | 1.92 |
| PCB 203 | 2.64 | 1.35 | PCB 203 | 0.75 |
| **ΣPCB** | **195.16** | **100.00** | **ΣPCB** | **100.01** |

**Table S7.** PCB congeners found in the serum of children from the urban area of East Chicago, IN (Marek et al., 2013). Congeners in red are impurities present in the neat PCB congeners used to prepare this mixture.

| **(A) Congers Detected above the LOQ in Serum** | | | **(B) Synthetic PCB Congener Profile** | |
|---|---|---|---|---|
| **Congener** | **Amount (ng/g lipid)** | **Percent of Total** | **Congener** | **Percent of Total** |
| PCB 18+30 | 3.3 | 3.18 | PCB 18+30 | 3.18 |
| PCB 20+28 | 7.79 | 7.51 | PCB 20+28 | 8.04 |
| PCB 21+33 | 3.02 | 2.91 | PCB 21+33 | 3.21 |
| PCB 31 | 4.97 | 4.79 | PCB 31 | 5.64 |
| PCB 56 | | | PCB 56 | 0.09 |
| PCB 61+70+74+76 | | | PCB 61+70+74+76 | 0.04 |
| PCB 66 | 4.55 | 4.39 | PCB 66 | 4.77 |
| PCB 83 | 11.77 | 11.34 | PCB 83 | 11.72 |
| PCB 84 | 3.02 | 2.91 | PCB 84 | 3.60 |
| PCB 85+116+117 | 3.75 | 3.61 | PCB 85+116+117 | 3.03 |
| PCB 90+101+113 | | | PCB 90+101+113 | 0.04 |
| PCB 105 | 4.51 | 4.35 | PCB 105 | 3.48 |
| PCB 110+115 | 9.49 | 9.15 | PCB 110+115 | 7.78 |
| PCB 118 | 13.45 | 12.96 | PCB 118 | 15.38 |
| PCB 129+137+138+164+164 | 11.86 | 11.43 | PCB 129+137+138+164+164 | 10.57 |
| PCB 147+149 | 2.65 | 2.55 | PCB 147+149 | 1.99 |
| PCB 153+168 | 9.95 | 9.59 | PCB 153+168 | 8.98 |
| PCB 156+157 | 1.57 | 1.52 | PCB 156+157 | 1.27 |
| PCB 177 | | | PCB 177 | 0.03 |
| PCB 180+193 | 5.62 | 5.42 | PCB 180+193 | 4.62 |
| PCB 181 | | | PCB 181 | 0.03 |
| PCB 182 | | | PCB 182 | 0.06 |
| PCB 184 | | | PCB 184 | 0.08 |
| PCB 187 | 2.5 | 2.41 | PCB 187 | 2.29 |
| PCB 188 | | | PCB 188 | 0.05 |
| PCB 209 | | | PCB 209 | 0.02 |
| **ΣPCB** | **103.77** | **100.00** | **ΣPCB** | **99.99** |

**Table S8.** PCB congeners found in the serum of mothers from the rural area of Columbus Junction, IA (Marek et al., 2013). Congeners in red are impurities present in the neat PCB congeners used to prepare this mixture.

| **(A) Congers Detected above the LOQ in Serum** | | | **(B) Synthetic PCB Congener Profile** | |
|---|---|---|---|---|
| **Congener** | **Amount (ng/g lipid)** | **Percent of Total** | **Congener** | **Percent of Total** |
| PCB 20+28 | 6.12 | 3.16 | PCB 20+28 | 3.01 |
| PCB 21+33 | 1.53 | 0.79 | PCB 21+33 | 2.04 |
| PCB 31 | 2.99 | 1.55 | PCB 31 | 1.74 |
| PCB 37 | 1.80 | 0.93 | PCB 37 | 0.72 |
| PCB 56 | | | PCB 56 | 0.04 |
| PCB 66 | 4.49 | 2.32 | PCB 66 | 2.23 |
| PCB 83 | 13.34 | 6.90 | PCB 83 | 6.71 |
| PCB 84 | 2.26 | 1.17 | PCB 84 | 1.22 |
| PCB 90+101+113 | | | PCB 90+101+113 | 0.05 |
| PCB 99 | 3.89 | 2.01 | PCB 99 | 1.92 |
| PCB 105 | 3.47 | 1.79 | PCB 105 | 2.13 |
| PCB 108 | | | PCB 108 | 0.12 |
| PCB 110+115 | 6.01 | 3.11 | PCB 110+115 | 4.57 |
| PCB 118 | 18.52 | 9.57 | PCB 118 | 12.55 |
| PCB 128+166 | | | PCB 128+166 | 0.12 |
| PCB 129+137+138+163+164 | 27.45 | 14.19 | PCB 129+137+138+163+164 | 16.61 |
| PCB 146 | 1.85 | 0.96 | PCB 146 | 1.00 |
| PCB 147+149 | | | PCB 147+149 | 0.10 |
| PCB 153+168 | 33.29 | 17.21 | PCB 153+168 | 8.10 |
| PCB 156+157 | 6.16 | 3.18 | PCB 156+157 | 4.65 |
| PCB 167 | | | PCB 167 | 0.07 |
| PCB 170 | 14.52 | 7.51 | PCB 170 | 5.41 |
| PCB 172 | | | PCB 172 | 0.06 |
| PCB 174 | | | PCB 174 | 0.03 |
| PCB 177 | | | PCB 177 | 0.05 |
| PCB 180+183 | 29.51 | 15.25 | PCB 180+183 | 11.99 |
| PCB 187 | 9.37 | 4.84 | PCB 187 | 9.78 |
| PCB 198+199 | 6.90 | 3.57 | PCB 198+199 | 2.97 |
| **ΣPCB** | **193.47** | **100.00** | **ΣPCB** | 99.99 |

**Table S9.** PCB congeners found in the serum of children from the rural area of Columbus Junction, IA (Marek et al., 2013). Congeners in red are impurities present in the neat PCB congeners used to prepare this mixture.

| (A) Congers Detected above the LOQ in Serum | | | (B) Synthetic PCB Congener Profile | |
|---|---|---|---|---|
| **Congener** | **Amount (ng/g lipid)** | **Percent of Total** | **Congener** | **Percent of Total** |
| PCB 8 | 2.42 | 2.90 | PCB 8 | 3.67 |
| PCB 18+30 | 1.97 | 2.36 | PCB 18+30 | 2.39 |
| PCB 20+28 | 5.88 | 7.05 | PCB 20+28 | 7.98 |
| PCB 31 | 4.52 | 5.42 | PCB 31 | 6.81 |
| PCB 37 | 2.44 | 2.93 | PCB 37 | 3.59 |
| PCB 83 | 8.85 | 10.62 | PCB 83 | 7.43 |
| PCB 84 | 3.07 | 3.68 | PCB 84 | 4.55 |
| PCB 90+101+113 | | | PCB 90+101+113 | 0.04 |
| PCB 105 | 1.87 | 2.24 | PCB 105 | 1.73 |
| PCB 108 | | | PCB 108 | 0.04 |
| PCB 110+115 | 7.34 | 8.81 | PCB 110 | 7.55 |
| PCB 118 | 10.66 | 12.79 | PCB 118 | 15.97 |
| PCB 129+137+138+163+164 | 12.51 | 15.01 | PCB 129+138+163 | 14.39 |
| PCB 153+168 | 11.76 | 14.11 | PCB 153/168 | 13.26 |
| PCB 167 | | | PCB 167 | 0.03 |
| PCB 177 | | | PCB 177 | 0.04 |
| PCB 180+193 | 10.07 | 12.08 | PCB 180+193 | 10.42 |
| PCB 181 | | | PCB 181 | 0.03 |
| PCB 182 | | | PCB 182 | 0.07 |
| PCB 209 | | | PCB 209 | 0.02 |
| **ΣPCB** | **83.36** | **100.00** | **ΣPCB** | **100.01** |

**Table S10.** Similarity coefficient (cos Θ) between the profiles of targeted PCB mixture and the lab synthesized mimic mixture. The values of cos Θ between 0 and 1, 0 indicates the two sets of data are with no relationship, and 1 indicate the two sets of data are identical.

| **PCB mixture** | **cos Θ** |
|---|---|
| Indoor air of new construction buildings | 0.9926 |
| Air of Schools in the Urban Area of East Chicago, IN | 0.9663 |
| Air of Schools from Columbus Junction, IA | 0.9889 |
| Tissue of Predatory Lake Fish from Cook County, IL. | 0.9904 |
| Tissue of Bottom Dwelling Lake Fish from Cook County, IL | 0.9873 |
| Serum of Mothers from the Urban Area of East Chicago, IN | 0.9370 |
| Serum of Children from the Urban Area of East Chicago, IN | 0.9923 |
| Serum of Mothers from the Rural Area of Columbus Junction, IA | 0.9263 |
| Serum of Children from the Rural Area of Columbus Junction, IA | 0.9853 |
